# Characterization of SARS-CoV-2 viral diversity within and across hosts

**DOI:** 10.1101/2020.05.07.083410

**Authors:** Palash Sashittal, Yunan Luo, Jian Peng, Mohammed El-Kebir

## Abstract

In light of the current COVID-19 pandemic, there is an urgent need to accurately infer the evolutionary and transmission history of the virus to inform real-time outbreak management, public health policies and mitigation strategies. Current phylogenetic and phylodynamic approaches typically use consensus sequences, essentially assuming the presence of a single viral strain per host. Here, we analyze 621 bulk RNA sequencing samples and 7,540 consensus sequences from COVID-19 patients, and identify multiple strains of the virus, SARS-CoV-2, in four major clades that are prevalent within and across hosts. In particular, we find evidence for (i) within-host diversity across phylogenetic clades, (ii) putative cases of recombination, multi-strain and/or superinfections as well as (iii) distinct strain profiles across geographical locations and time. Our findings and algorithms will facilitate more detailed evolutionary analyses and contact tracing that specifically account for within-host viral diversity in the ongoing COVID-19 pandemic as well as future pandemics.

## INTRODUCTION

The current COVID-19 outbreak, caused by a novel coronavirus, severe acute respiratory syndrome coronavirus 2 (SARS-CoV-2), has become a global pandemic and still keeps spreading, despite extensive mitigation efforts. As of May 7^th^, 3.78 million COVID-19 cases have been confirmed in more than 100 countries, with 269,000 fatalities. A deep understanding of the virus’ evolution and transmission patterns is critical to enable effective outbreak control and mitigation strategies. In contrast to the majority of existing studies, that have characterized viral genomes based on consensus sequences, we look into the heterogeneity within individual samples and shared mutational signatures across samples. It is critical to evaluate viral diversity below the consensus level as minor variants may impact the patterns of virulence and person-to-person transmission efficiency. Here, we report a genomic analysis of 621 SARS-CoV-2 samples with high-quality sequencing data and 7,540 samples from the GISAID database with consensus sequences. Although recent studies have looked at within-host diversity of COVID-19 patients (Rose et al., 2020; Ramazzotti et al., 2020; Shen et al., 2020; Karamitros et al., 2020; Tang et al., 2020), this study is, to the best of our knowledge, the first comprehensive analysis at a large scale that attempts to identify genomic signatures of SARS-CoV-2 strains that occur within and across individual hosts. Through genomic signature deconvolution, we identify multiple strains that are present in the global population and find evidence of co-existence of distinct strains within hosts.

We analyzed 621 bulk sequencing samples, that met stringent quality control criteria, from the Sequence Read Archive (SRA) database deposited before April 22^nd^, 2020. At the time of developing the algorithms for our analysis, we only had access to samples deposited before April 9^th^, 2020. As such, we considered a discovery set of 161 high-quality Illumina samples deposited before April 9^th^, 2020 and a validation set with the remaining 460 samples deposited before April 22^nd^, 2020 (Figure 1A). We developed algorithms to deconvolve the bulk sequencing samples in the discovery set and to subsequently find the proportion of the inferred strains in each SRA sample. The inferred strains were validated against 7,540 GISAID consensus sequences deposited before April 15^th^, showing that our strains cover a large fraction (94%) of this database.

**Figure 1:**
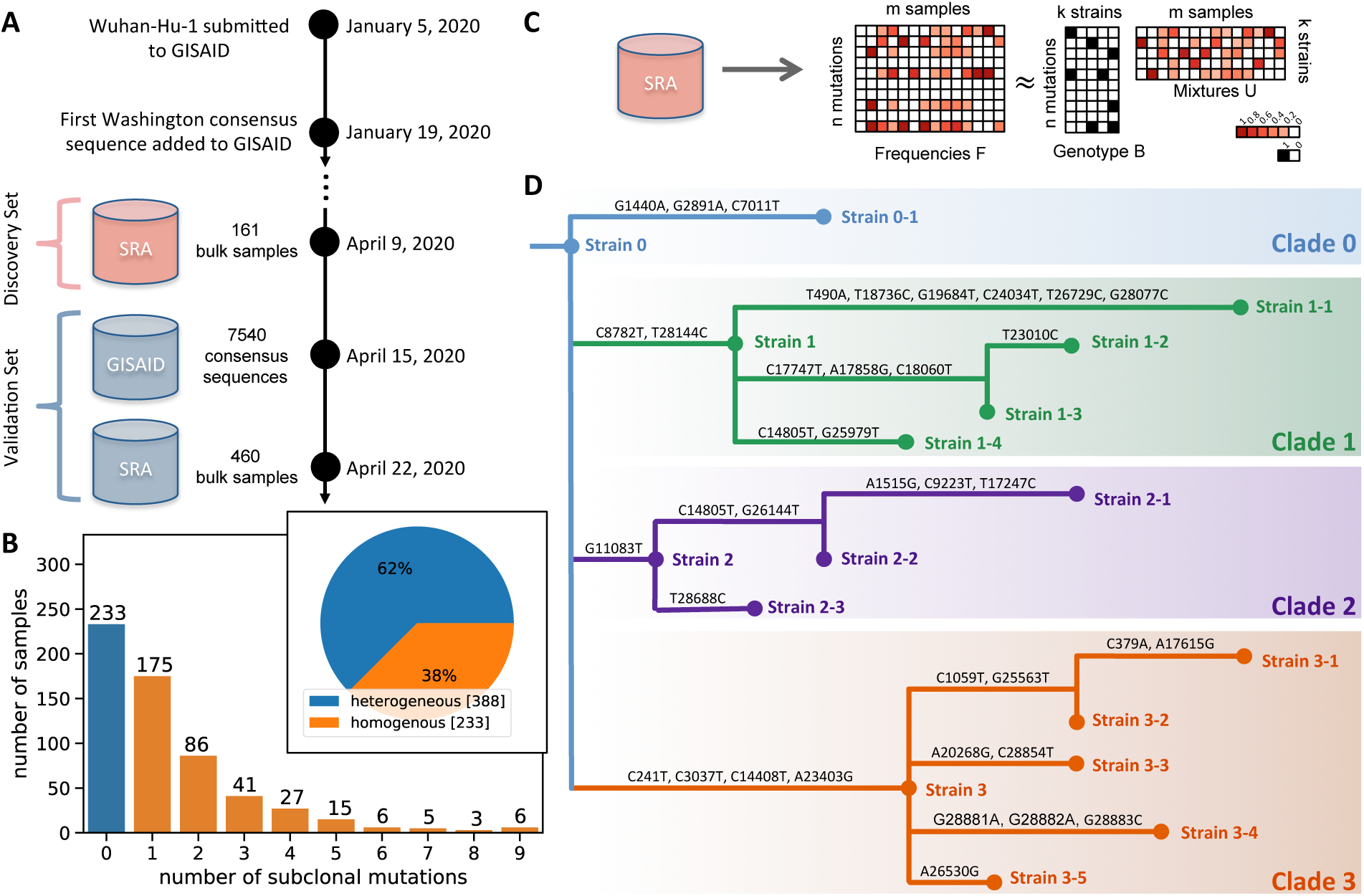
Deconvolution Framework to Identify SARS-CoV-2 Viral Strains Within and Across Hosts. (A) Timeline showing the collection dates of the discovery (161 high-quality Illumina samples) and validation set (460 Illumina/nanopore samples and 7,540 GISAID consensus sequences) used in the study. (B) The majority of sequencing samples (388/621) contain subclonal mutations, indicative of within-host SARS-CoV-2 viral diversity. The frequency of samples with varying number of subclonal mutations is also shown. (C) In the Strain Deconvolution problem, we are given the variant allele frequency (VAF) matrix *F*, containing the VAF of every mutation in each sample, and a number *k* of strains to be inferred. Our goal is to infer the genotype matrix *B* and mixture matrix *U* such that *F* ≈ *BU*, thus elucidating strains that occur within and across COVID-19 hosts along with their sample-specific proportions. (D) The phylogenetic tree on the 17 strains inferred using the discovery set and validated against the GISAID sequences in the validation set. We identify four distinct clades in the tree (colors) with distinct characteristics in terms of geographical and temporal spread, transmittability and mutability. See also Figures S1 and S2.

Following a phylogenetic analysis of the inferred strains, we clustered the strains into four distinct clades (Figure 1D), which match the phylogenetic tree inferred on the GISAID consensus sequences (Figure 2C). We performed a spatiotemporal analysis and found that although Clade 3 arose most recently (Figure S1E), it is the most prevalent clade in most North American and European countries (Figure 2D). Separate epidemiological analyses showed that strains from Clade 3 have a higher average reproduction number compared to the other clades (Figure S3). At the protein-level we found that the missense mutations considered in this study occur primarily on protein surfaces and are non-deleterious (Figure 3A). Finally, our deconvolution approach enabled us to look at viral diversity below the consensus level, identifying hosts with strains from distinct clades as well as multiple cases of recombination (Figure 4). Both these phenomena hint at multistrain infections and/or superinfections. These findings have important implications for our understanding of the transmissibility and mutability of specific strains of SARS-CoV-2. In particular, examining the viral composition of COVID-19 patients below the consensus level will enable more precise contact tracing.

**Figure 2:**
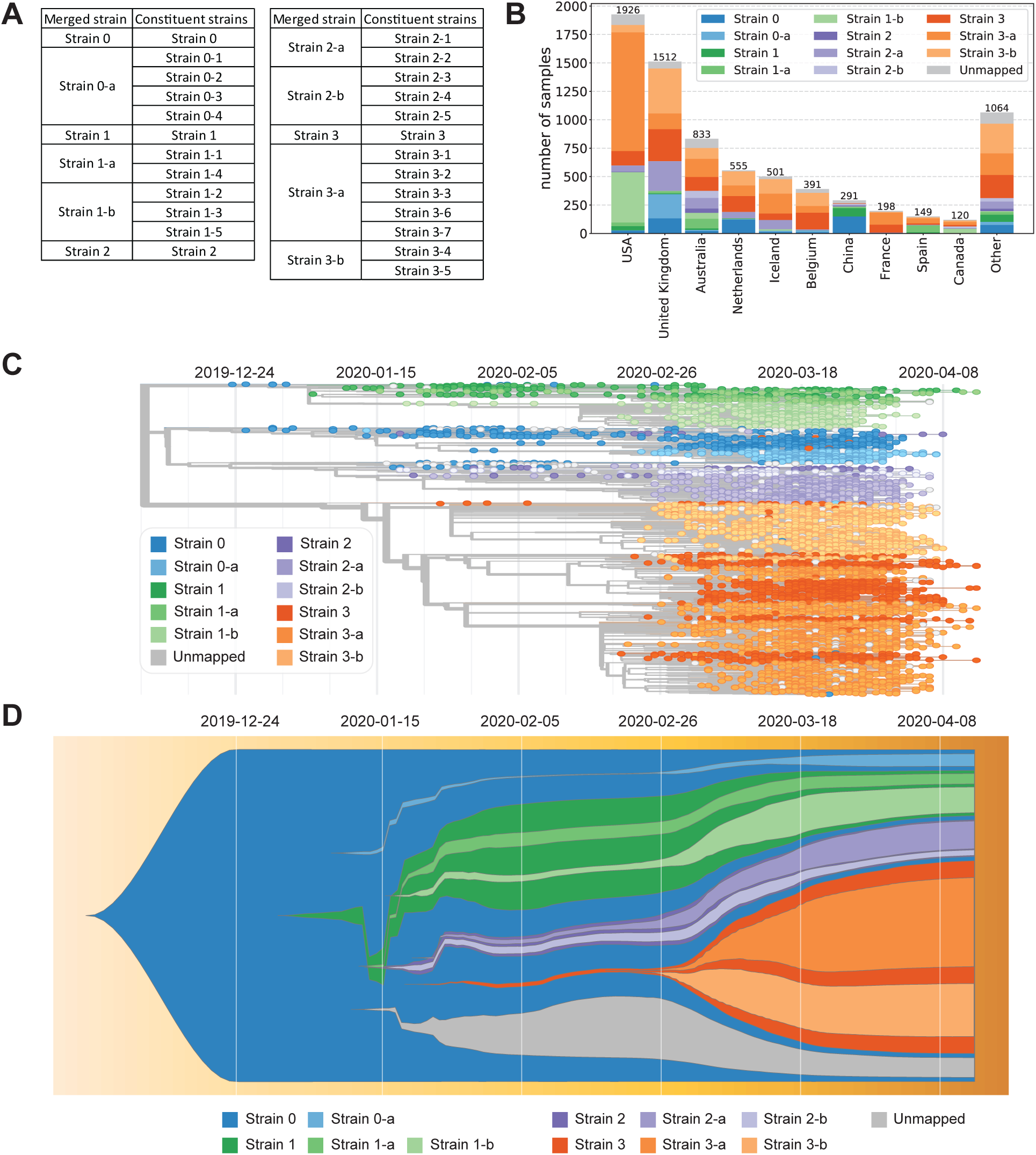
Evolutionary, Temporal and Geographical Distribution of Identified SARS-CoV-2 Strains. (A) Grouping of identified *k* = 25 strains used in (B), (C) and (D). (A) Number of GISAID consensus sequences from each country and how they map to our identified strains. (C) The phylogenetic tree for all 7,540 GISAID sequences. Each sample (leaf in the tree) is colored by the strain to which it maps. (D) This fishplot shows the proportion of GISAID sequences that map to each strain as a function of time (increasing from left to right). See also Figure S3.

**Figure 3:**
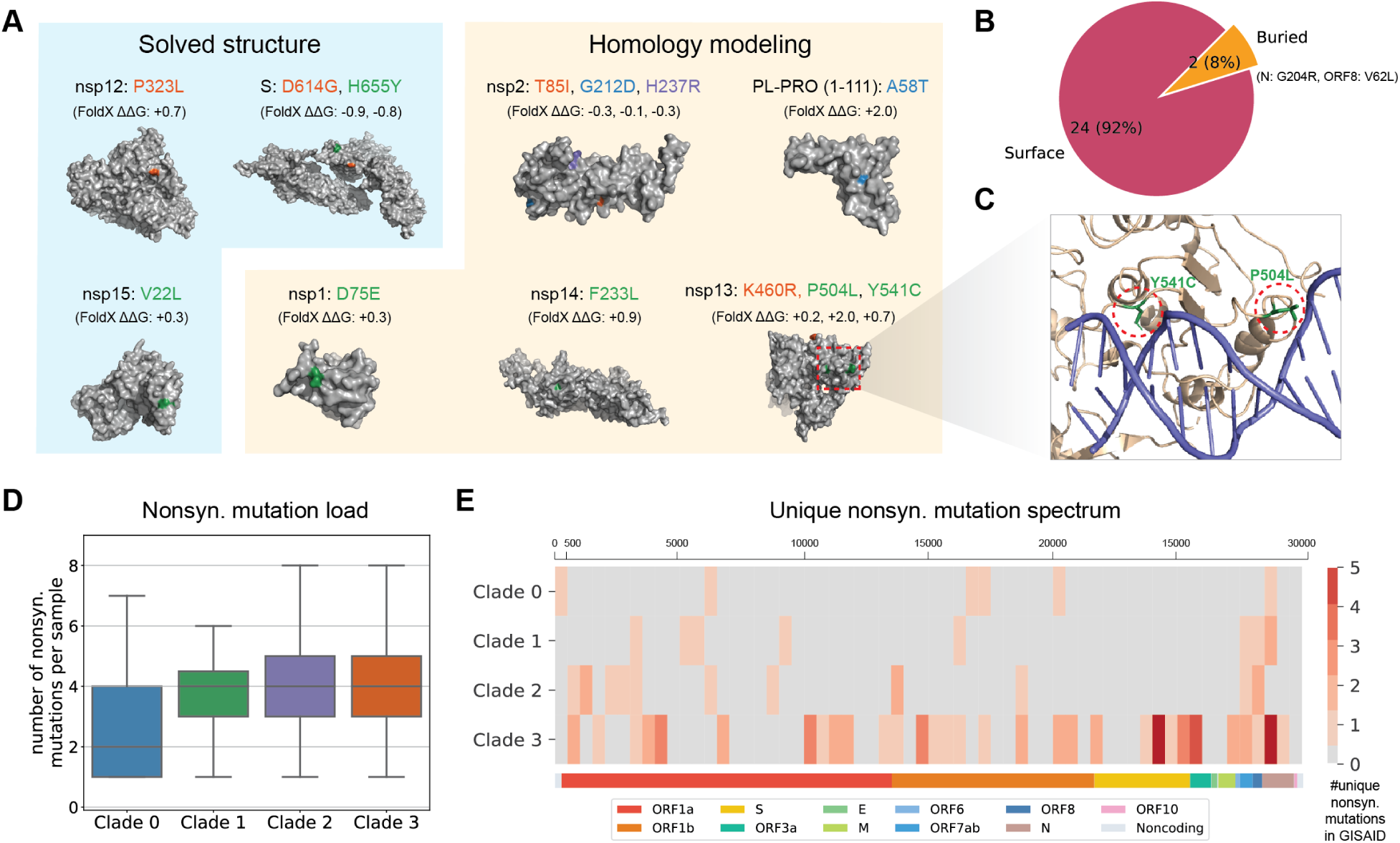
Identified Missense Mutations are mostly on Surface and Non-deleterious. (A) Among the 43 identified mutations, a subset of 28 are missense, 13 of which are present on solved structures and high-quality homology models. Structure stability changes (ΔΔ*G*) inferred using FoldX (Delgado et al., 2019) were shown in Kcal/mol. Mutations are colored according to their clade. (B) 24/26 mutations are surface facing while 2/26 are buried. (C) A zoom-in view of the docking results of nsp13 and double-stranded nucleic acid. Two mutations, P504L and Y541C, are highlighted. (D) Box plots of nonsynonymous mutation load, showing the number of nonsynonymous mutations per GISAID sample of each clade. (E) Unique nonsynonymous mutation spectrum of the SARS-CoV-2 genome. For each clade and a bin size of 500bp on the genome, the number of unique nonsynonymous mutations GISAID samples is counted. Only recurrent nonsynonymous mutations (using a threshold of ten samples) are visualized in the heatmap. See also Figure S4.

**Figure 4:**
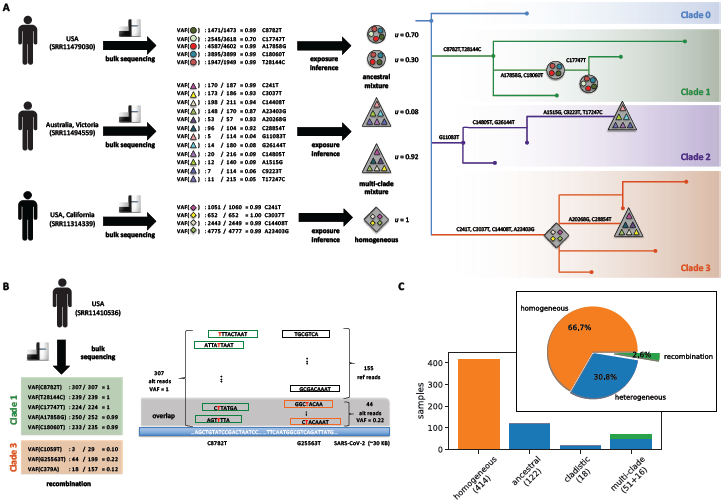
Deconvolution Reveals Cases of Within-host Viral Diversity, Recombination and Multi-strain Infection. (A) We show three example samples with clonal and subclonal mutations and the inferred strains present within them. The position of the inferred strains in the phylogenetic tree is also shown. We show one homogeneous samples and two heterogeneous samples, out of which one is an *ancestral mixture* and the other is a *multi-clade mixture*. (B) We show the example of a sample that has a strong evidence for recombination between strains from Clade 1 and Clade 3. The sample has 5 mutations from Clade 1 and 3 mutations from Clade 3. An alternative explanation with no recombination event would require parallel evolution of Strain 1-3 with the three mutations from Clade 3. (C) The within-host diversity of the 621 SRA samples is characterized into four classes – *homogeneous, ancestral mixture, cladistic mixture* and *multi-clade mixture*. 66.7% of the samples are homogeneous while the rest are heterogeneous. Moreover, 2.6% of the samples show some evidence of recombination between strains belonging to distinct clades. See also Figure S5.

## RESULTS

### Identification of SARS-CoV-2 Genomic Signatures

We analyzed 621 SARS-CoV-2 sequencing samples from the NCBI Sequence Read Archive (SRA) to identify SARS-CoV-2 genomic signatures/strains that occur within and across COVID-19 patients (Figure 1A). These samples were selected among a larger pool of 1,217 SRA samples, satisfying stringent filtering criteria regarding depth and breadth of coverage (STAR Methods). We used a standard bioinformatics pipeline to align the reads of each sequencing sample to the SARS-CoV-2 reference genome followed by removal of duplicate reads and single-nucleotide variant calling with stringent quality criteria (STAR methods). We define a mutation as *subclonal* if it has a variant allele frequency (VAF) between 0.05 and 0.95 and is supported by at least 5 variant reads. Importantly, a subclonal mutation is only present in a subsets of viral strains within a host and is thus indicative of the presence of multiple strains within that host. On the other hand, a *clonal mutation*, i.e. a mutation supported by at least 5 variants and a VAF of at least 0.95, is present among all strains within a host. Strikingly, we found that 388/621 samples (62%) contain subclonal mutations and thus are *heterogeneous*, with 213/621 samples (34%) containing two or more subclonal mutations (Figure 1B).

To study whether these subclonal mutations are present on recurring strains, we devised a deconvolution algorithm. The underlying mathematical problem, Strain Deconvolution, is a variant of the classical nonnegative matrix factorization problem (Paatero and Tapper, 1994) (STAR Methods). That is, given a matrix *F* ∈ [0,1]^*n*×*m*^ of variant allele frequencies of *n* mutations across *m* samples, we seek a mutation-by-strain genotype matrix *B* ∈ {0,1}^*n*×*k*^ and a strain-by-sample mixture matrix *U* ∈ [0,1]^*k*×*m*^ such that *F* ≈ *BU* and the proportions of each strain sum to 1 (Figure 1C). We solve this problem by decomposition into two subproblems that we alternately solve until convergence. Using simulations, we validated that this algorithm convergences to the ground truth solution (STAR Methods).

To prevent overfitting and spurious mutations due to sequencing artifacts, we split the bulk sequencing samples into a *discovery set* composed of 161 high-quality Illumina samples and a *validation set* composed of the remaining 460 Oxford nanopore and Illumina samples. To begin, we focus our attention on the discovery set, which is comprised of samples from Australia, USA, China, Germany and Nepal. Upon solving the Strain Deconvolution problem for the discovery set, we identified *k* = 25 strains (Figure S2) composed of *n* = 43 mutations (Table S2). We built a phylogeny for the *k* = 25 identified strains, resulting in four major clades (Figure 1D and Figure S2). The number of strains per clade varied from 5 to 8. Examining the collection dates of samples in the discovery set, we find that Clade 0 is comprised of the earliest samples (January 2020) whereas Clade 3 corresponds to the most recent samples (Figure S1E).

### Evolutionary and Geographical Distribution of Identified Viral Strains

Next, we corroborated the *k* = 25 identified viral strains using 7,540 GISAID consensus sequences in the validation set (Figure 1A). To do so, we categorized a GISAID consensus sequence as belonging to a strain if the sequence was identical to that strain (or one of its ancestors – see STAR Methods) in terms of our *n* = 43 mutations. For visualization purposes we grouped strains together as described in Figure 2A. We found that our *k* = 25 strains covered 7,096 sequences (94%) in GISAID with varying proportions across countries (Figure 2B). In particular, we found that Australia appears to be very diverse in terms of strain composition, whereas the USA seems to primarily contain strains from Clade 3. On the other hand, China seems to contain strains mainly from Clade 0 and Clade 1. In particular, all strains from Clade 1 contain mutations C8782T and T28144C, which was previously termed the ‘L-strain’ (Tang et al., 2020).

Futhermore, we found that the phylogeny we constructed from our identified *k* = 25 strains (Figure 1D and Figure S2) is consistent with the Nextstrain phylogeny inferred from GISAID sequences, with matching clades and strains (Figure 2C). Figure 2D shows a fishplot (Miller et al., 2016) that highlights the temporal dynamics of relative strain prevalences across the globe, indicating the dominance of strains from Clade 3. Inspection of this figure shows that Strains 1-2, 1-3 and 1-5 (grouped together as Strain 1-b) quickly rose to dominance within Clade 1. These strains share common mutations in nsp13 that we will characterize below. We see a similar pattern for Strains 2-1 and 2-2 (grouped together as Strain 2-a) that have become more prevalent than Strains 2-3, 2-4 and 2-5 (grouped together as Strain 2-b).

To more precisely investigate the transmission dynamics of our strains, we computed the average *R*_0_ value of the strains from each clade from March 1^st^, 2020 to April 13^th^, 2020 (STAR Methods). In line with Figure 2D, we found that strains from Clade 3 have a higher *R*_0_ value of 3.60 compared to a mean of 1.93, 1.99 and 2.32 of other strains from Clade 0, Clade 1 and Clade 2, respectively (Figure S3). One possible explanation for this difference is that all the strains within Clade 3 have a nonsynonymous mutation A23403G, orresponding to a D614G amino acid substitution in the S (spike) protein, that has been hypothesized to increase transmissibility (Korber et al., 2020).

Finally, we performed an immunogenicity analysis, showing a weak relationship between the clade composition and the HLA allele composition. We first applied MHCflurry (O’Donnell et al., 2018) to compute the HLA/MHC-I peptide specificity of all mutational positions and counted the number of strong binders with predicted IC50 ≤ 500 nM for each HLA allele. Then we approximated the immunogenicity of a sample by taking the average of the count of binders for all HLA allele types weighted by the HLA frequencies of the country where the sample is from (Gonzalez-Galarza et al., 2019). We performed the immunogenicity comparison at the clade level in four major countries (USA, UK, Australia and the Netherlands) with the most GISAID sequences and focused on the clades with at least 50 samples. In the USA, Clade 1 and Clade 3 seem to have fewer binders than Clade 0 and Clade 2, which is correlated with the prevalence of the clades. Similar trends can be found also in the UK, Australia, and Netherlands (Figure S4D).

### Characterization of Nonsynonymous Mutations in Genomic Signatures

Among the 43 genomic mutations in the strain signatures, 28 of them are nonsynonymous, including 27 missense mutations which cause amino acid changes and one nonsense mutation (ORF7a-E95*) which yields a truncated protein product (Table S2). To investigate the impact of missense mutations on protein structure, we searched the protein databank (PDB) (Berman et al., 2000) and found three solved structures: nsp12 (Yin et al., 2020), nsp15 (Kim et al., 2020) and the S protein (Walls et al., 2020). We then used HHpred (Söding et al., 2005) to build homology models for the remaining SARS-CoV-2 proteins. Among these, we were able to identify high-confidence homologous templates for nsp1, nsp2, nsp13, nsp14 and a domain of PL-PRO (1-111). For the remaining proteins, we collected publicly available protein structure predictions by a variety of protein folding algorithms. In particular, we obtained the predicted structures for the M protein [Feig-lab’s refined RaptorX model (Heo and Feig, 2020; Källberg et al., 2012)], N protein [Zhang lab’s C-I-TASSER model (Zhang et al., 2020)], nsp6 [Feig-lab’s refined AlphaFold model (Heo and Feig, 2020; Jumper et al., 2020; Senior et al., 2020)], ORF3a [Feig-lab’s refined RaptorX model (Heo and Feig, 2020; Källberg et al., 2012)], ORF8 [Feig lab’s model (Heo and Feig, 2020)], and PL-PRO [1260-1945, Feig-lab’s refined RaptorX model (Heo and Feig, 2020; Källberg et al., 2012)]. We were able to map 26 missense mutations to these structures, except one mutation on the S protein (R682W) which is not solved. Figure 3A shows mutations on solved structures and high-quality homology models. We found that 24 out of 26 mutations are near or on the surface of these proteins (Figure 3B). The impact of 25 surface mutations on the stability of protein structures, calculated with the FoldX forcefield (Delgado et al., 2019), are very small (i.e. ΔΔ*G <* 3 Kcal/mol), indicating that these mutations are not deleterious and that these amino acid changes with little impact on the protein structure were likely to be transmitted. Among the mutations on solved structures or high-quality models, we found that Strain 1-3’s signature includes two mutations on the same protein nsp13 (P504L and Y541C). This protein is a helicase and leads the replication and transcription complex to unwind duplex RNA and DNA, in addition to its function in viral self-reproduction (Yu et al., 2012; Jia et al., 2019). To investigate the role of the two mutations in nucleic acid binding, we took the homology model of SARS-CoV-2 nsp13, which has 100% amino-acid sequence identity with the SARS-CoV nsp13 structure template, and performed docking with double-stranded DNA and single-stranded DNA. Strikingly, although the two surface mutations, P504L and Y541C, are not in direct contact (C-*β* distance ≈ 19 Angstrom), they are likely to jointly coordinate the binding to double-stranded nucleic acid as well as single-stranded nucleic acid (Figure 3 and Figure S4D). These results are consistent with the hydrogen/deuterium exchange differences and electrophoretic mobility shift assay performed on SARS-CoV nsp13 (Jia et al., 2019).

Beyond the signature mutations, we also stratified the nonsynonymous mutations that appeared in the GISAID samples according to their clade classification (Figure 3D and Figure 3E). We found that Clade 1 and 2 have similar mutation loads, while Clade 0 has a lower load, and Clade 3 has a slightly higher load, both are statistically significant (p-value = 2.37 × 10^−51^ and 3.36 × 10^−7^, respectively) (Figure 3). This is consistent with the prevalence and the evolutionary history of these four clades. Further, we looked at the number of distinct nonsynonymous mutations that occur in more than 10 GISAID samples that map to a given clade. Clade 3 has a higher number of such unique nonsynonymous mutations than the other three clades, covering most proteins with a few specific hotspots on S (spike) protein and ORF3a (Figure 3E). When comparing the mutation loads between synonymous and nonsynonymous mutations, we noticed that Clade 3 appears to have higher nonsynonymous loads and lower synonymous loads compared to Clade 1 and Clade 2, suggesting that Clade 3 may evolve under a higher level of positive selection (Figure S4E).

### Evidence for Within-host Diversity, Recombination and Multi-strain Infection

While Figure 1C demonstrates that a large fraction (62%) of SARS-CoV-2 sequencing samples exhibit within-host diversity, it does not enable us to conclude which of our *k* = 25 strains are present within a host. To that end, we posed the Strain Exposure problem, where given a genotype matrix *B* and variant allele frequencies **f** of a sequencing sample we seek to identify which strains are present in that sample along with their proportions, imposing an additional sparsity constraint to prevent overfitting (STAR Methods). In addition, we considered an augmented matrix *B* which includes ancestral strains composed of subsets of the mutations along the branches of the phylogenetic tree (Figure S2). We formulated the Strain Exposure problem as a mixed integer quadratic program that we solved using Gurobi (Gurobi Optimization, 2020). We solved this problem for the 612 SRA samples in the discovery and validation set.

We classified a sample as exhibiting within-host diversity (*heterogeneous*) if at least two distinct strains are inferred to be present in that sample. We found that 207/602 samples satisfy this criterion (Figure 4C). Note that this number of samples is smaller than that inferred in Figure 1D as in this case we restricted our analysis to the *n* = 43 mutations that define our *k* = 25 strains (Table S2). If a sample only has a single strain, we classified it as *homogeneous* (67% of samples). We further classified the within-host diversity of each sample into three categories based on the phylogenetic relationships among the strains inferred to be present in the sample. The first category is *ancestral mixture*, which means that there exists a path from the root to some node in the phylogenetic tree in Figure S2 such that all the strains present in the sample lie on this path. We found that 122 samples (20%) can be categorized as *ancestral mixtures*. The second category is *cladistic mixture*, which means that all the strains belong to the same clade but the sample is not an *ancestral mixture*. This happens when the sample has strains from different branches in the same clade. We found that 18 samples (3%) fall into this category. Finally, the third category is *multi-clade mixture*, which has 67 samples (11%). Samples that belong to this category contain at least two strains from distinct clades.

Figure 4A describes three example samples – the clonal and subclonal mutations in the samples, the inferred strains and their position in the phylogenetic tree shown in Figure 1B. Sample SRR11314339 originates from California, USA and has four mutations, all of which are clonal. Our algorithm inferred that there exists only one strain (Strain 3) in this sample and therefore this sample is classified as *homogeneous*. The second sample we show is SRR11494559 from Victoria, Australia. This sample has 12 mutations of varied variant allele frequencies. We inferred that these mutations and their frequencies can be explained by the presence of two strains, Strain 2-1 and Strain 3-3, each with 6 mutations. Since these strains belong to distinct clades, namely Clade 2 and Clade 3, we categorized this sample as a *multi-clade mixture*. The third example sample SRR11479030 is from the USA and has five mutations, of which only one is subclonal. The four clonal mutations have VAF almost equal to 1, which means that any strain present in this sample must contain these four mutations. The presence of the one subclonal mutation (C17747T) in this sample is explained by the presence of a descendant strain that contains all five of the mutations. We identified the strain that contains these five mutations as Strain 1-3 from Figure S1D. Since this sample only contains Strain 1-3 and its ancestor, we classified it as *ancestral mixture*.

Among the 67 samples that belong to the *multi-clade mixture* category, we found 16 samples that show evidence of putative recombination. These samples have at least two mutations from strains belonging to distinct clades such that the sum of their VAFs is more than 1. When this happens, there must be a strain in the sample that harbors both mutations. This is a direct consequence of the pigeon-hole principle. We observed this phenomenon in 16 SRA samples. In addition to recombination, there are several other explanations for the presence of these mutations in the same strain such as parallel evolution, back mutations or sequencing errors. We can rule out sequencing errors due to our stringent criteria for selecting the mutations (see STAR Methods). If the presence of many mutations in a sample need to be attributed to homoplasy (parallel evolution or back mutations) then a single recombination event is a more parsimonious explanation. We considered a sample as having strong evidence of putative recombination if the number of such mutations in the sample is three or more. An example of such a sample, SRR11410536 from the USA, is shown in Figure 4B. This sample has mutations from two separate clades – five mutations from Clade 1 and three mutations from Clade 3. Due to the high VAF values of these mutations (close to 1 for mutations from Clade 1 and more than 0.1 for mutations from Clade 3), we inferred that at least one strain present in this sample must contain all these mutations concurrently. The strain that contains all the mutations from Clade 1 present in this sample is Strain 1-3. One possible scenario is that this sample contains a descendant of Strain 1-3 that has the three additional mutations from Clade 3 due to parallel evolution. However, a more parsimonious explanation would be a single recombination event between Strain 1-3 and a strain that contains the three mutations from Clade 3 present in this sample, such as Strain 3-1 or Strain 3-2. A total of 8 SRA samples show similarly strong evidence of putative recombination (Table S1).

Viral diversity within a host can arise due to within-host evolution after infection by a single strain, or a single infection event composed of multiple genetically-distinct strains (*multi-strain infections*) or multiple, separate infection events (*superinfections*). Figure S5A shows that there are 19 mutations (out of 43 mutations considered in our analysis) that are present subclonally in more than 5 SRA samples. While the shared subclonal mutations could have arisen independently in multiple samples due to parallel evolution, a more parsimonious explanation is the transmission of multiple strains through multi-strain infections and/or superinfections. This concept was first introduced in cancer genomics to test for the presence of complex patterns of seeding in metastasis, such as polyclonal seeding and re-seeding (Brastianos et al., 2015; ElKebir et al., 2018). Indeed, our method inferred the co-occurrence of multiple strains in the SRA samples, shown in Figure S5E, which include strains that have high support in the GISAID samples (Figure 2). This observation, together with the 67 *multi-clade mixture* and *recombination* samples, point towards multi-strain and/or superinfections occurring in the COVID-19 pandemic.

## DISCUSSION

In this work, we performed the first, comprehensive large-scale study of SARS-CoV-2 viral diversity within and across COVID-19 hosts. We analyzed 621 bulk-sequencing samples from the Sequence Read Archive (SRA) and 7,540 consensus sequences from the GISAID database (Elbe and Buckland-Merrett, 2017). Specifically, we developed a deconvolution algorithm that we applied to a smaller discovery set of 161 SRA samples. We obtained *k* = 25 strains defined by *n* = 43 mutations that are distributed across four major clades with distinct patterns of geographical and temporal spread. In particular, Clade 3, has a potentially higher transmissibility rate, a higher level of nonsynonymous mutational load, and a broader mutational spectrum compared to the other clades.

Among the analyzed 621 samples, 207 samples contained multiple strains, with a subset of 67 samples containing strains from distinct clades. In addition, we found evidence of putative recombination between strains from distinct clades in 16 samples. The most parsimonious explanation for both these findings is multi-strain and/or superinfection. Our protein-level analysis showed that all the missense mutations considered in our study were non-deleterious and mostly near or on the protein surface. In particular, we found a pair of potentially compensatory missense mutations in nsp13 that may play a conserved role in the nucleic acid binding function of this helicase protein.

There are several avenues for future research. First, we expect that our algorithms can be applied to analyze SARS-CoV-2 viral diversity at the organ level. Second, correlating our identified strains with outcomes such as severity, mortality and morbidity is of great importance to further understand the pathogenesis of COVID-19. Third, it will be interesting to study whether the presence of distinct strains within a host has adverse outcomes. Fourth, the strains inferred for each infected host can improve the reconstruction of the transmission history using methods that support multiple samples per host and multi-strain infections (De Maio et al., 2018; Sashittal and El-Kebir, 2020). Finally, we expect more precise contact tracing when one takes within-host diversity into account.

**Table 1:** Identified Mutations and their Phylogenetic Clades for the Tree Shown in Figure 1D. For each mutation, we list its gene name, protein name, amino acid (AA) change, structure availability, and clade membership. The “Struct. avail.” column indicates the sources of the protein structures we collected: proteins that have solved structures in PDB are labeled with “solved” and the PDB IDs are provided; other proteins are labeled with “homology” if high-confidence homology models can be built; predicted structures are collected from folding algorithms for the remaining proteins, which are labeled with “folding” (STAR Methods); “-” means the mutation is either in a noncoding region or a synonymous/nonsense mutation, or there is no available structure that models the mutated position. See also Table S2 for the list of mutations in the extended phylogenetic tree shown in Figure S2.

| SNV | Gene | Protein | AA | Struct. avail. | Clade 0 | Clade 1 | Clade 2 | Clade 3 |
| --- | --- | --- | --- | --- | --- | --- | --- | --- |
| C241T | noncoding |  | - | - | - | - | - | ✓ |
| C379A | ORF1a | nsp1 | V38V | - | - | - | - | ✓ |
| T490A | ORF1a | nsp1 | D75E | homology | - | ✓ | - | - |
| C1059T | ORF1a | nsp2 | T85I | homology | - | - | - | ✓ |
| G1440A | ORF1a | nsp2 | G212D | homology | ✓ | - | - | - |
| A1515G | ORF1a | nsp2 | H237R | homology | - | - | ✓ | - |
| G2891A | ORF1a | PL-PRO | A58T | homology | ✓ | - | - | - |
| C3037T | ORF1a | PL-PRO | F106F | - | - | - | - | ✓ |
| C7011T | ORF1a | PL-PRO | A1431V | folding | ✓ | - | - | - |
| C8782T | ORF1a | nsp4 | S76S | - | - | ✓ | - | - |
| C9223T | ORF1a | nsp4 | H223H | - | - | - | ✓ | - |
| G11083T | ORF1a | nsp6 | L37F | folding | - | - | ✓ | - |
| C14408T | ORF1b | nsp12 | P323L | solved (PDB:7BV1) | - | - | - | ✓ |
| C14805T | ORF1b | nsp12 | Y455Y | - | - | ✓ | ✓ | - |
| T17247C | ORF1b | nsp13 | R337R | - | - | - | ✓ | - |
| A17615G | ORF1b | nsp13 | K460R | homology | - | - | - | ✓ |
| C17747T | ORF1b | nsp13 | P504L | homology | - | ✓ | - | - |
| A17858G | ORF1b | nsp13 | Y541C | homology | - | ✓ | - | - |
| C18060T | ORF1b | nsp14 | L7L | - | - | ✓ | - | - |
| T18736C | ORF1b | nsp14 | F233L | homology | - | ✓ | - | - |
| G19684T | ORF1b | nsp15 | V22L | solved (PDB:6VWW) | - | ✓ | - | - |
| A20268G | ORF1b | nsp15 | L216L | - | - | - | - | ✓ |
| T23010C | S | S | V483A | folding | - | ✓ | - | - |
| A23403G | S | S | D614G | solved (PDB:6VXX) | - | - | - | ✓ |
| C24034T | S | S | N824N | - | - | ✓ | - | - |
| G25563T | ORF3a | ORF3a | Q57H | folding | - | - | - | ✓ |
| G25979T | ORF3a | ORF3a | G196V | folding | - | ✓ | - | - |
| G26144T | ORF3a | ORF3a | G251V | folding | - | - | ✓ | - |
| A26530G | M | M | D3G | folding | - | - | - | ✓ |
| T26729C | M | M | A69A | - | - | ✓ | - | - |
| G28077C | ORF8 | ORF8 | V62L | folding | - | ✓ | - | - |
| T28144C | ORF8 | ORF8 | L84S | folding | - | ✓ | - | - |
| T28688C | N | N | L139L | - | - | - | ✓ | - |
| C28854T | N | N | S194L | folding | - | - | - | ✓ |
| G28881A | N | N | R203K | folding | - | - | - | ✓ |
| G28882A | N | N | R203R | - | - | - | - | ✓ |
| G28883C | N | N | G204R | folding | - | - | - | ✓ |

## Acknowledgments

We thank all the authors who have kindly deposited and shared genome data on GI-SAID (https://www.gisaid.org). A full table of acknowledgments for GISAID contributors can be found here: https://github.com/elkebir-group/SARS-CoV-2-deconvolution/master/data/metadata.tsv. This material is based upon work supported by the National Science Foundation under award numbers DBI-1652815, CCF-1850502 and CCF-2027669.

## Author Contributions

All authors conceived the methodology. P.S. implemented the bioinformatics pipeline. P.S. and M.E.-K. implemented the strain deconvolution and exposure inference algorithm. P.S., J.P. and M.E.-K. performed the strain deconvolution validations and analyzed the results. P.S. performed the epidemiological analysis. Y.L. and J.P. performed the protein analyses. Y.L. visualized the evolutionary, temporal and geographical distributions of the GISAID sequences. J.P. and M.E.-K. supervised the research. All authors wrote the paper.

## Key Resource Table

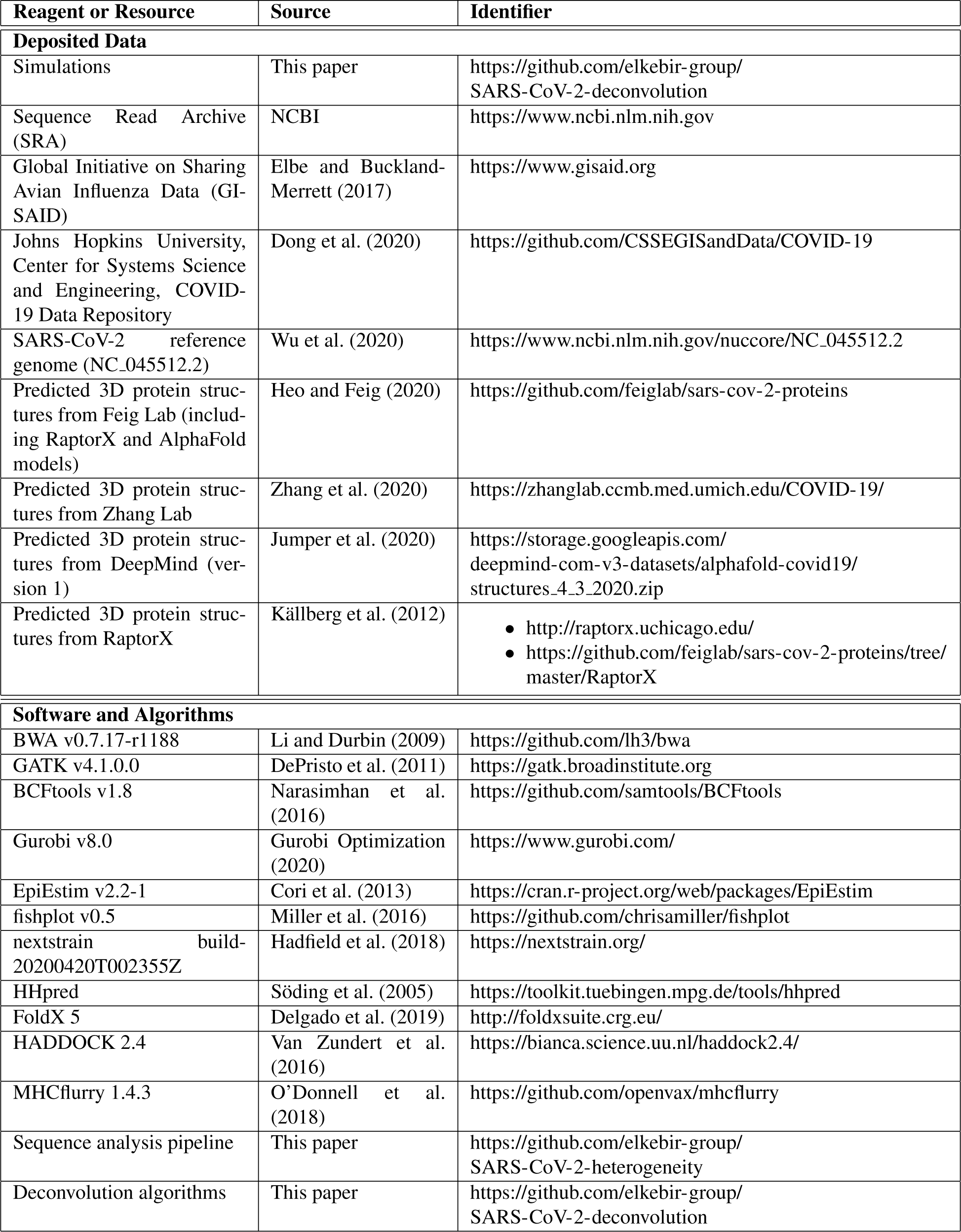

## STAR⋆METHODS

### LEAD CONTACT AND MATERIALS AVAILABILITY

Further information and requests for resources and reagents should be directed to and will be fulfilled by the Lead Contact, Mohammed El-Kebir. This study did not generate any new materials.

## METHOD DETAILS

### Bulk Sequencing Data

We downloaded 1,217 SARS-CoV-2 sequencing samples from the SRA database deposited before April 22^nd^, 2020. We used a bioinformatics pipeline to (i) align reads from each sample to the SARS-CoV-2 reference genome, (ii) remove duplicate reads and (iii) identify single-nucleotide variants (SNVs). Specifically, we used BWA (Li and Durbin, 2009) to map sequencing reads to the SARS-CoV-2 reference genome (NC 045512.2). Subsequently, we marked PCR duplicates using GATK (DePristo et al., 2011). For SNV calling, we only considered samples that have a mean depth (number of reads supporting a position in the genome) of 50 and coverage breadth (number of positions in the genome mapped by at least one read) of 20%. We called the SNVs using bcftools (Narasimhan et al., 2016), requiring a Phred score of at least 20 and mapping quality also at least 20. We used the Illumina sequencing samples deposited before April 9^th^, 2020 as the discovery dataset to identify strains that occur within and across hosts by solving the Strain Deconvolution problem. Specifically, we imposed the following criteria for a sample to be included in the discovery set (i) it must contain at least one mutation that is also present in another sample in the discovery set, (ii) it must have at least one subclonal mutation (i.e. its variant allele frequency greater than 0.05 and less than 0.95) and (iii) it must be from a unique host. After we removed samples that do not pass these criteria, 161 SRA samples with SNVs at 97 positions were kept as the discovery set for the identification of SARS-CoV-2 viral strains. The validation set is comprised of 460 SRA samples deposited in the SRA database before April 22^nd^. The selection criteria for these samples was that they must have a mean depth of 50 at the 43 positions that were selected after we perform the Strain Deconvolution on the discovery dataset (described in the following section).

### Strain Deconvolution via Nonnegatve Matrix Factorization

We formulated the problem of strain deconvolution as a specific nonnegative matrix factorization. Consider a sequencing sample obtained from a host infected by one or more strains of SARS-CoV-2. After alignment and quality control, as described above, we consider *n* positions in the viral genome for deconvolution, all of which also exhibit exactly two alleles, the reference allele and the alternate allele, and assume that *k* strains exist within and across the samples in the discovery dataset. We define the *genomic signature* of a strain as a binary vector **b** = [*b*_*i*_] ∈ {0,1}^*n*^, where *b*_*i*_ = 1 if the strain contains the variant allele at the *i*-th position and *b*_*i*_ = 0 otherwise. Thus, the genotypes of all *k* strains in the sample are given by the *genotype matrix B* = [*b*_*i,j*_] ∈ {0,1}^*n*×*k*^. The proportions of *k* strains in the sample are given by the mixture vector **u** = [*u*_*j*_] such that *u*_*j*_ ≥ 0 for each strain *j* and 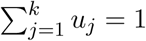. For each sample in the discovery set, we computed the mutation frequencies **f** = [*f*_*i*_] ∈ [0,1]^*n*^ which may be caused by mixtures or recombinations of the strains in the sample. More specifically, *f*_*i*_ denotes the frequency of the variant allele at position *i* in the sample. Since we only have the observed frequencies, we expect to solve the following problem. Given a frequency vector **f**, we want to find a genotype matrix *B* and mixture vector **u** such that **f** can be reconstructed or approximated by *B***u**. For the entire discovery dataset with *m* ≥ 1 samples from different infected hosts, with a frequency matrix *F* = [*f*_*i,p*_] ∈ [0,1]^*n*×*m*^, we optimize an *n* × *k* genotype matrix *B* and *k* × *m* mixture matrix *U* to reconstruct the given frequency matrix *F*, formulated as follows.

#### Problem 1 (Strain Deconvolution).

*Given frequency matrix F* ∈ [0,1]^*n*×*m*^ *and number k of strains, find a genotype matrix B* ∈ {0,1}^*n*×*k*^ *and mixture matrix U* ∈ [0,1]^*k*×*m*^ *such that (i)* **1**^*T*^ *U* = **1**^*T*^ *and (ii)* 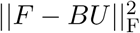 *is minimum*.

Similar matrix decomposition problems arise in several other biological applications such as gene network inference (Liao et al., 2003), mutational signature inference in cancers (Alexandrov et al., 2013) and deconvolving cell-mixtures from DNA methylation data (Houseman et al., 2012). An efficient algorithm to solve the Strain Deconvolution problem in a low-rank setting has been shown in Slawski et al. (2013). In this study, we approximately solved this problem using a penalty method approach (Bach et al., 2012) by relaxing the binary constraints on *B*. A penalty function is introduced to gradually enforce the *B* matrix to become binary. Specifically, we solved the following relaxed optimization problem

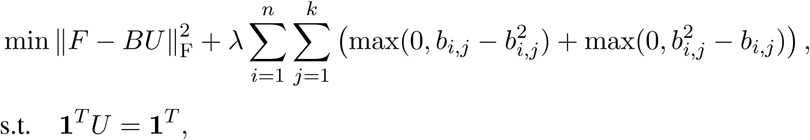

where *U* ∈ [0,1]^*k*×*m*^ and *B* ∈ ℝ^*n*×*k*^. The first term in the objective function is the *reconstruction error*, and the second term is the *penalty function*, which becomes zero when *b*_*i,j*_ is binary, and is otherwise positive. For entries in *F* matrix with a missing value, we omit the corresponding errors when computing the reconstruction loss. We solve this problem with a two-stage iterative algorithm. Specifically, in each iteration, we alternately minimize the objective function with respect to *B* while fixing *U*, and then solve for *U* while fixing *B*. T he p roblem o f o ptimizing *B* given *U* is solved using a Limited-memory Broyden–Fletcher–Goldfarb–Shanno (L-BFGS) algorithm (Liu and Nocedal, 1989). We formulated the second problem of optimizing *U* given *B* as a quadratic program with linear constraints and solve it using Gurobi. Initially, we set *λ* = 1 in the first iteration *t* = 1 and increase by a factor of 1 .005 in every subsequent iteration to strengthen the penalty. We stopped increasing *λ* when *λ* ≥ 4. We also stabilize the optimization using the dampening trick by mixing (*U*^*t*−1^,*B*^*t*−1^) from the previous iteration *t* − 1 and the solution (*U,B*) obtained in the current iteration *t*, yielding final matrices (*U*^*t*^,*B*^*t*^) upon completing the *t*−th iteration defined as

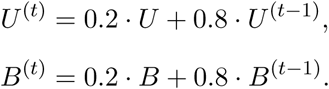

The convergence is achieved when the *reconstruction error* is less than a pre-defined threshold a maximum number of iterations has been reached.

### Validation of our Deconvolution Algorithm using Simulations

We assessed our deconvolution approach using simulated data, where ground truth is known. Ground truth consists of a genotype matrix *B* ∈ {0,1}^*n*×*k*^ and a mixture matrix *U* ∈ [0,1]^*k*×*m*^. Here, we consider *n* = 100 mutations, *m* = 50 samples and *k* ∈ {5,10} strains. In addition, we vary the fraction *m*_*f*_ ∈ {0, 0.1} of missing entries. Specifically, we generate a genotype matrix *B* by setting each entry *b*_*ij*_ = 1 with probability 3/*n* resulting in an expected number of 3 mutations per strain. The mixture matrix *U* is generated by drawing the number *k*_*p*_ of strains per sample *p* ∈ [*m*] from a zero-truncated Poisson distribution with mean 3. Subsequently, *k*_*p*_ strains are drawn without replacement and assigned mixture proportions by a draw from a symmetric Dirichlet distribution with the concentration parameter equal to 1. For each combination of *n, m, k* and *m*_*f*_, we generated 20 simulated instances, composed of a frequency matrix *F* and ground truth solution (*B,U,k*). This amounts to a total of 80 simulation instances.

We ran our deconvolution algorithm for 100 iterations and 20 restarts, choosing for each simulation instance the solution that minimizes the objective function 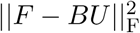 among the restarts. Figure S1A shows that our method achieves very small normalized reconstruction error. As such, our method recovers the simulated genotype matrix in all cases, as quantified by the Hamming distance shown in Figure S1B.

### Application of Deconvolution Algorithm to Identify SARS-CoV-2 Viral Strains

We performed the deconvolution on our discovery set (the 161 SRA samples with SNVs at 97 positions). To determine *k*, we first performed the singular value decomposition of the matrix *F* ∈ [0,1]^*n*×*m*^. The missing entries are replaced with zero and the singular values are arranged in descending order of magnitude. We saw a sharp drop in the singular values with a rank 25 approximation of *F* having a Frobenius norm that is 98.5% of the Frobenius norm of *F* (Figure S1C), which suggested the low-rank structure of *F*. We then ran a sweep on the number of strains *k* ∈ {5,10,15,20,25,30,35,40,45,50} and solve the Strain Deconvolution problem for each *k* independently with 25 random initializations until convergence. Figure S1D shows the convergence of our method for increasing number of iterations. The improvement of reconstruction error started to be saturated at *k* = 25 or 30. To be parsimonious, we selected *k* = 25 and identified 25 strains with 43 of the 97 mutations covered. We checked the rest mutations which were not included and found that they tend to have very low variant allele frequencies and most of them only appeared in two samples. Therefore, our method identifies the SNVs that are shared across samples while also inferring the underlying strains.

### Phylogenetic Analysis of Identified SARS-CoV-2 Viral Strains

We used RAxML (Stamatakis, 2014) from the Augur (Neher and Bedford, 2015) pipeline to generate the phylogenetic tree over the *k* = 25 viral strains. The phylogenetic tree with four identified clades is shown in Figure S2. The ancestral strains, placement of mutations on branches and the amino-acid changes were inferred using Augur (Neher and Bedford, 2015). We used the open-sourced software Nextstrain and its visualization tool Auspice (Hadfield et al., 2018) to visualize the evolutionary distribution in Figure 2C. Coloring and labeling schemes in Nextstrain were adapted to visualize our identified strains. We used the ‘fishplot’ R package (Miller et al., 2016) to visualize the temporal distribution of the identified strains in Figure 2D.

### Identification of Ancestral Strains

We used the phylogenetic tree (Figure S2) to enumerate all possible ancestral strains on each branch of the phylogeny. We assumed that mutations that have occurred in some strain cannot be lost in its descendant strains. Using this approach we obtain a total of 129 different strains. Supplementary Data 1 contains the exact mutational makeup of each ancestral strain. The clade distribution of the 129 ancestral strains is as follows

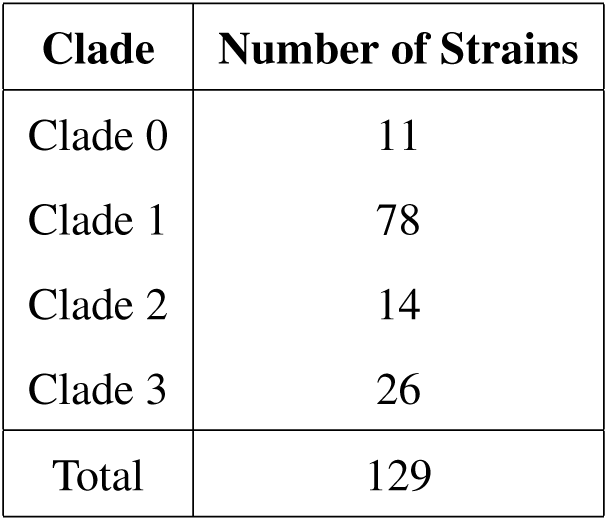

### Strain Specific Epidemiological Analysis

In order to understand the transmissibility of the identified strains of SARS-CoV-2, we used EpiEstim (Cori et al., 2013; Thompson et al., 2019) version 2.2-1 to infer the reproduction number *R*_0_(*t*) for each clade as a function of time *t*. For this analysis, we assumed that strains belonging to distinct clades are transmitted independently, which is in line with the observation that only the minority of SRA samples consists of strains from distinct clades (67 out 621). We obtained the number of confirmed cases of COVID-19 for each day after 22^nd^ January from the “COVID-19 Data Repository by the Center for Systems Science and Engineering (CSSE) at Johns Hopkins University” (Dong et al., 2020). These cases are annotated by the country of origin as well. In order to infer the reproduction number for a given clade, we need to find the number of confirmed cases infected with strains of that clade for each day. We approximated this by multiplying the number of confirmed cases on a given day by the proportion of GISAID sequences mapped to some strain of a given clade on that day. The underlying assumption here is that the GISAID sequences have been uniformly sampled from the infected population. Therefore, to avoid sampling errors, we restricted our analysis to 6 countries (Australia, Netherlands, Spain, China, USA and United Kingdom) that have at least 2 clades such that more than 50 GISAID sequences map to a strain from those clades.

Briefly, EpiEstim Cori et al. (2013) models transmission as a Poisson process where the rate of infection depends on the reproduction number *R*_0_(*t*) at time *t* and the infectivity of the infected individual. The infectivity of an infected individual is assumed to only depend on the time elapsed since infection, *i*.*e*. the infectivity of a person after time *s* since infection is given by *w*(*s*). The function *w* is known as the infectivity distribution. The reproduction number *R*_0_(*t*) at any time *t* is estimated by the ratio of the number of new infections to the total infectiousness of the infected individuals at time *t*. The total infectiousness at any time is highly dependent on the infectivity distribution of the disease. In practice it is approximated by distribution of the serial interval, which is the time between the onset of symptoms in a primary case and the onset of symptoms in secondary cases. In our study, following the work of Cori et al. (2013) and Menendez (2020), we assume the serial distribution to be a Gamma distribution with mean *µ*_*SI*_ and standard deviation *σ*_*SI*_. Since we do not know the exact values of *µ*_*SI*_ and *σ*_*SI*_ for the COVID-19 outbreak, we draw these values from normal distributions with the constraint that *µ*_*SI*_ *> σ*_*SI*_. In a recent study of 94 patients with laboratory-confirmed COVID-19, the mean of the serial interval distribution was estimated to be 5.8 days (He et al., 2020). Another study based on 28 infector-infectee pairs of COVID-19 patients estimated the mean serial interval to be 4.6 days (Nishiura et al., 2020). Based on these results, we draw the mean *µ*_*SI*_ from a normal distribution with mean 5 days and standard deviation of 1 day. The standard deviation *σ*_*SI*_ is drawn from a normal distribution with mean 2.9 days and standard deviation 1 day. We used 200 sample pairs of (*µ*_*SI*_, *σ*_*SI*_) in our study. The posterior distribution of time varying reproduction number *R*_*t*_ is formed by 200 realization of *R*_*t*_ for each pair of (*µ*_*SI*_, *σ*_*SI*_). At each time *t*, we calculated the average reproduction number over a time-window of 7 days ending at time *t*.

Figure S3A shows the clade-specific reproduction number for each country separately. For each country, we only included the clades if more than 50 GISAID consensus sequences from the country perfectly map to a strain of that clade. The shaded region around each line shows the 95% credibility interval. We see that for all countries, except United Kingdom, the reproduction number of Clade 3 is higher than the other clades. The same trend is seen in Figure S3B that shows the clade-specific reproduction number aggregated over all the major countries shown in Figure S3A.

### Exposure Inference to Identify Strain Composition in COVID-19 Samples

Here we expect to identify the strain composition in COVID-19 samples with sequencing data. For a sample with a given frequency vector **f** ∈ [0,1]^*n*^ and the genotype matrix *B* ∈ [0,1]^*n*×*k*^ of a set *S* of *k* strains, we expect to find a smaller number of strains from *S* such that (i) each selected strain is present in a sufficient proportion in the sample, and (ii) the deconvolution error for the sample defined by 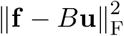 is minimized. To this end, we solved the Strain Exposure problem defined as follows.

#### Problem 2

(Strain Exposure). *Given a frequency vector* **f** ∈ [0,1]^*n*^, *a genotype matrix B* ∈ {0,1}^*n*×*k*^, *number ℓ of strains and thresholds τ* ∈ [0,1], *find a mixture vector* **u** ∈ [0,1]^*k*^ *such that (i)* **1**^*T*^ **u** = 1, *(ii)* **u** *has at most C nonzero entries that are at least τ and the remaining entries are* 0, *and (iii)* 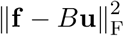 *is minimum*.

We formulated this problem as a mixed integer quadratic program and used Gurobi (Gurobi Optimization, 2020) to solve for increasing values of *ℓ* until the *reconstruction error* stops decreasing. The value of *τ* is set to 0.05, consistent with the definition of subclonal mutations.

### Classifying Within-host Diversity

Upon solving the Strain Exposure problem for SRA samples in the discovery and validation set, we classify the samples with within-host diversity into ancestral, cladistic, or multi-clade mixtures. The main text contains precise definitions of these three categories. For the latter category, multi-clade mixture, we further annotated samples as having possibly undergone recombination by considering the number of mutations that occur in strains from distinct clades. Table S1 lists the 16 SRA samples that show evidence of putative recombination.

### Immunogenicity Analysis

We used MHCflurry (O’Donnell et al., 2018) to predict MHC-peptide binding. We extracted all peptides of length 9 (9-mer) that cover any of the amino acid substitution in the list of our identified mutations (Table S2). We enumerated all the possible positions of the mutated residue within a 9-mer. The peptides containing the wildtype amino acid were also extracted for prediction. A peptide was considered a binder if the predicted affinity is below the standard IC50 cut-off of 500nM. We selected the allele-specific model in MHCflurry to predict affinity for samples in a country based on the HLA allele frequencies in that country. Specifically, we downloaded population HLA allele frequencies of countries from the Allele Frequency Net Database (Gonzalez-Galarza et al., 2019). If an allele has several frequencies available from different studies, we took the mean value as its frequency. We only predicted the affinity for alleles with frequency ≥ 5% in the country being considered.

### Protein Structure and Mutation Analysis

To characterize the 27 missense mutations on protein structure, we searched on the protein databank [PDB, Berman et al. (2000)] and found the solved structure for three SARS-CoV-2 proteins, including nsp12 [PDB ID: 7BV1, (Yin et al., 2020)], nsp15 [PDB ID: 6VWW, (Kim et al., 2020)] and S protein [PDB ID: 6VXX, (Walls et al., 2020)]. We then used HHpred (Söding et al., 2005) to build homology models and found high-confidence homologous templates for nsp1, nsp2, nsp13, nsp14 and a domain of PL-PRO (1-111). For the remaining proteins, we collected publicly available predicted protein structures by a variety of protein folding algorithms and obtained predicted structures for M protein [Feig-lab’s refined RaptorX model (Heo and Feig, 2020; Källberg et al., 2012)], N protein [Zhang lab’s C-I-TASSER model (Zhang et al., 2020)], nsp6 [Feig-lab’s refined AlphaFold model (Heo and Feig, 2020; Jumper et al., 2020; Senior et al., 2020)], ORF3a [Feig-lab’s refined RaptorX model (Heo and Feig, 2020; Källberg et al., 2012)], ORF8 [Feig lab’s model (Heo and Feig, 2020)], and PL-PRO [1260-1945, Feig-lab’s refined RaptorX model (Heo and Feig, 2020; Källberg et al., 2012)]. Among all the 27 missense mutations, we were able to map 26 of them to the structures we collected, except the S-R682W mutation that cannot be mapped to the solved region.

### Stability analyses

We performed the stability analysis of the missense mutations on protein structure using the FoldX forcefield (Delgado et al., 2019). We used the ‘RepairPDB’ function to refine collected protein structures and the ‘BuildModel’ function to obtain the change of stability (ΔΔ*G*) by taking the average of five independent runs.

### Protein-DNA Docking

The docking analyses of protein nsp13 that harbors two missense mutations were carried out using the HADDOCK 2.4 server (Van Zundert et al., 2016). The structure of nsp13 was obtained from the homology modeling using HHpred (Söding et al., 2005). We collected the structures of both double-strained DNA (dsDNA) and single-strainded DNA (ssDNA) from PDB (PDB IDs 4KNY and 2P6R, respectively). The HADDOCK scores, which measure the overall energy of the docking results, were also collected.

## DATA AND CODE AVAILABILITY

Our sequence analysis pipeline is available at https://github.com/elkebir-group/SARS-CoV-2-heterogeneity and the source code of our deconvolution algorithms can be downloaded from https://github.com/elkebir-group/SARS-CoV-2-deconvolution.

---

**Table S1:** 16 SRA Samples Show Evidence of Putative Recombination. Shown samples contain two mutations from distinct strains in distinct clades such that the sum of their VAFs exceeds 1. As a null hypothesis, we show the number of parallel evolution events required to explain the sample in the absence of recombination. We consider a sample to have strong evidence of recombination if the number of such parallel evolution events is more than or equal to 3.

| Sample Type | Identifier | Required Parallel Evolution Events |
| --- | --- | --- |
| validation | SRR11178053 | 2 |
| validation | SRR11178053 | $\geq 3$ |
| validation | SRR11178053 | 1 |
| validation | SRR11476467 | 2 |
| validation | SRR11476467 | 1 |
| discovery | SRR11410536 | $\geq 3$ |
| discovery | SRR11479040 | 1 |
| discovery | SRR11491746 | $\geq 3$ |
| discovery | SRR11491771 | $\geq 3$ |
| discovery | SRR11494523 | $\geq 3$ |
| discovery | SRR11494559 | $\geq 3$ |
| discovery | SRR11494585 | $\geq 3$ |
| discovery | SRR11494603 | 1 |
| discovery | SRR11494651 | 1 |
| discovery | SRR11494743 | $\geq 3$ |
| discovery | SRR11494753 | 1 |

**Figure S1:**
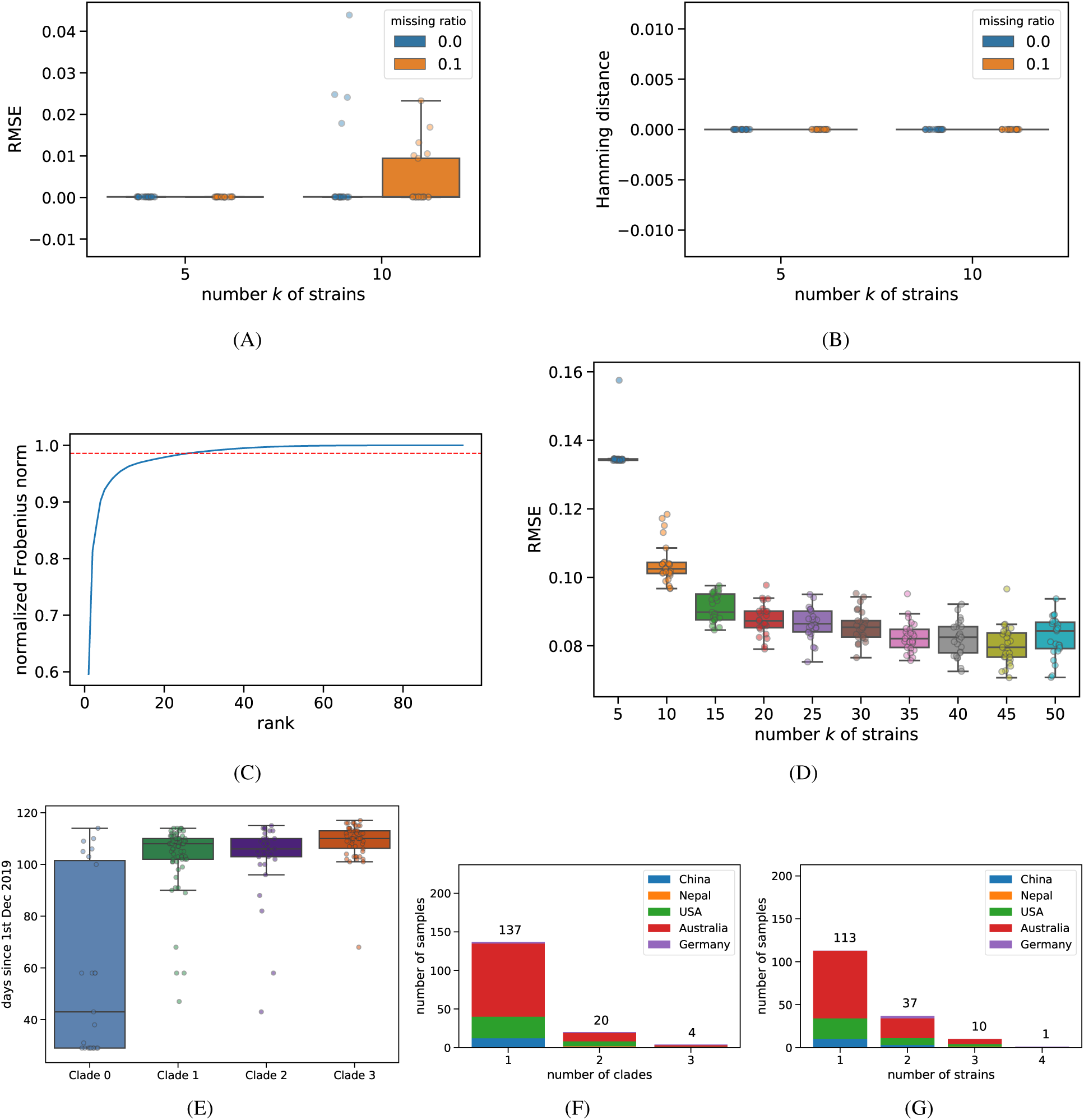
Validation of Deconvolution Approach via Simulations and Application to Discovery Data. (A) Root mean squared error (RMSE) for the simulated cases with number of strains *k* ∈ {5,10} and ratio *m*_*f*_ ∈ {0, 0.1} of missing entries. (B) The Hamming distance between the strains inferred by our method and the ground-truth strains. (C) The normalized Frobenius norm of the reduced-rank approximation of the variant allele frequency matrix *F* for the filtered discovery set comprising of 161 samples. The red dashed line shows the Frobenius norm for a rank 25 approximation of *F*. (D) RMSE of the deconvolution of the discovery data for each restart with increasing number *k* of strains. (E) For each clade, we show the collection dates of the discovery set samples with strains from that clade in the inferred solution. (F) Number of discovery set samples with strains that belong to increasing number of distinct clades in the inferred solution. (G) Number of discovery set samples with increasing number of strains in the inferred solution. Related to Figure 1.

**Figure S2:**
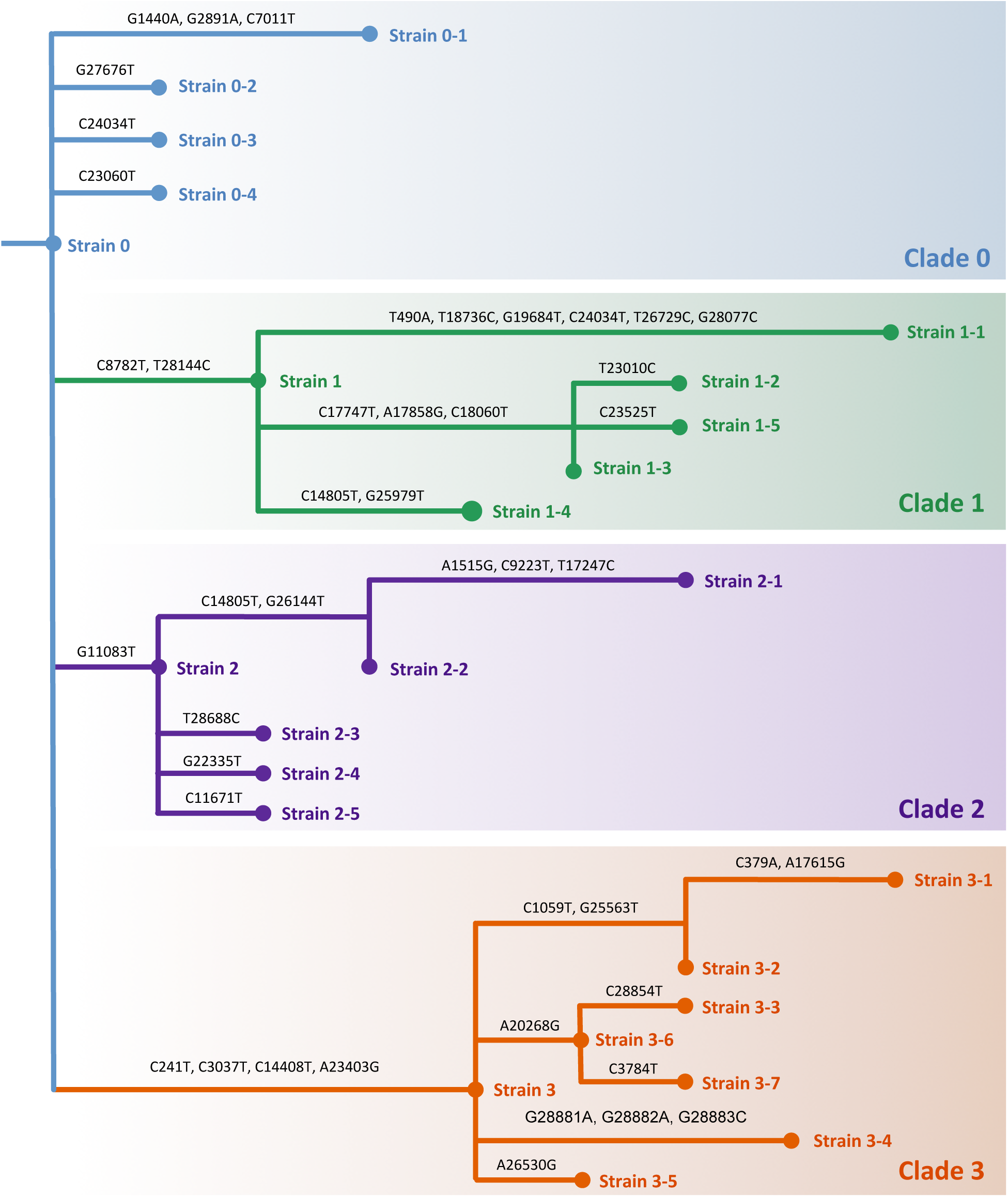
Phylogenetic Tree of the *k* = 25 Strains Inferred by Deconvolution of the Discovery Set. We identify four distinct clades in the tree (colors) with distinct characteristics. The branches are annotated by the nucleotide mutations introduced in the genome. The only positions that show homoplasy are 14805 (present in both Clade 1 and Clade 2) and 24034 (present in both Clade 0 and Clade 1). A reduced form of this tree with just 17 strains that have high support in the GISAID sequences is shown in Figure 2. Related to Figures 1 and 2.

**Figure S3:**
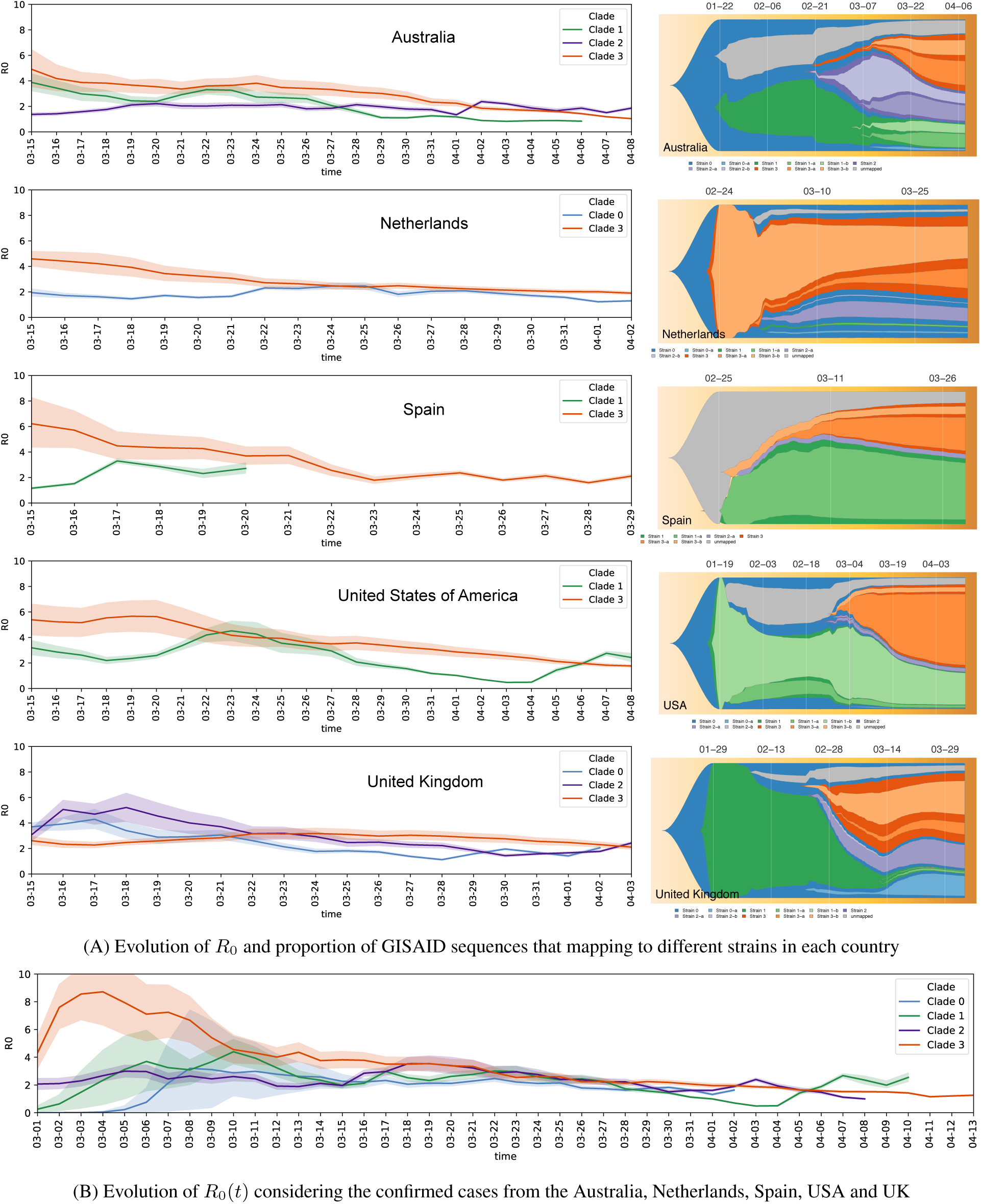
Transmissibility of Clade 3 is Higher than other Clades Despite its Recent Introduction. (A) The evolution of the reproduction number *R*_0_(*t*) for each clade in different countries in shown on the left. Clade 3 has the most transmissible strains in all the countries except United Kingdom. On the right we show the proportion of samples that map to different strains as the pandemic progresses in each country. Clade 3 is introduced the latest in all the countries except Netherlands. (B) The evolution of the reproduction number *R*_0_(*t*) for the total cases from all the countries considered in (A). Related to Figure 2.

**Figure S4:**
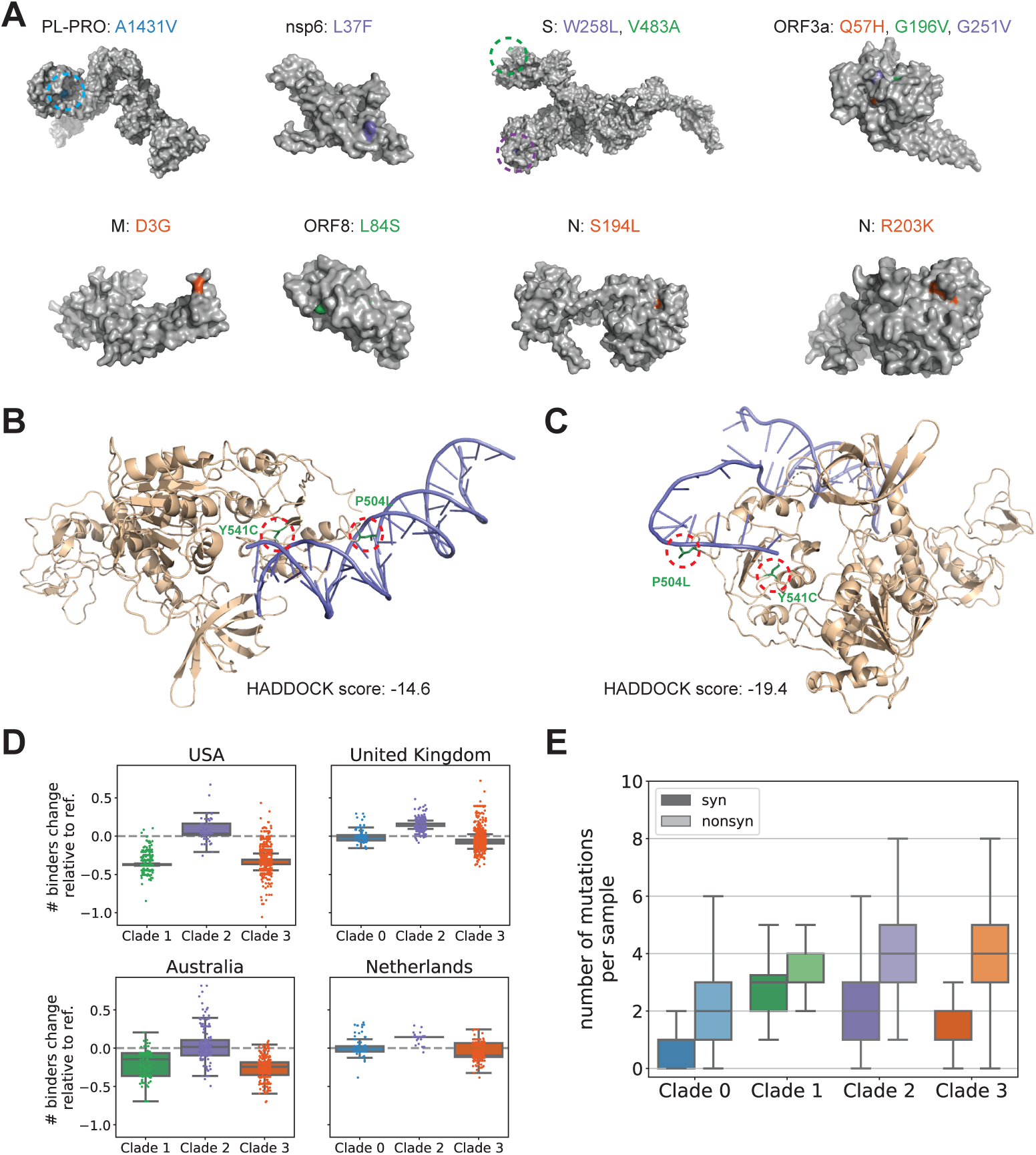
Most Nonsynonymous Mutations Considered in this Study are on Protein Surfaces. (A) In addition to the nonsynonymous mutations mapped to homology modelings or solved structures in Figure 3, we map the remaining 13 nonsynonymous mutations to public predicted protein structures from a variety of protein folding algorithms. Among these 13 mutations, 11 of them are on or near the protein surfaces while other two mutations, N-G204R and ORF8-V62L are buried. (B)-(C) Docking analyses are performed using HADDOCK 2.4 of nsp13, which harbors mutations P504L and Y541C, with (B) double-stranded DNA (dsDNA) and (C) single-stranded DNA (ssDNA), respectively. The two mutations are highlighted and the HADDOCK scores of the docking results, which measures the overall energy of the docking structure, are also shown. (D) Changes in the number of MHC binders (peptides binding to MHC) in samples from USA, United Kingdom, Australia, and Netherlands. Binders are counted as the weighted average based on the HLA allele frequencies of each country (STAR Methods). Each dot shows the difference of the number of binders between a GISAID sample and the reference sample are computed. Clades with less than 50 samples in a country are excluded from the analysis. (E) The number of synonymous (syn) and nonsynonymous (nonsyn) mutations per GISAID sample. Related to Figure 3.

**Figure S5:**
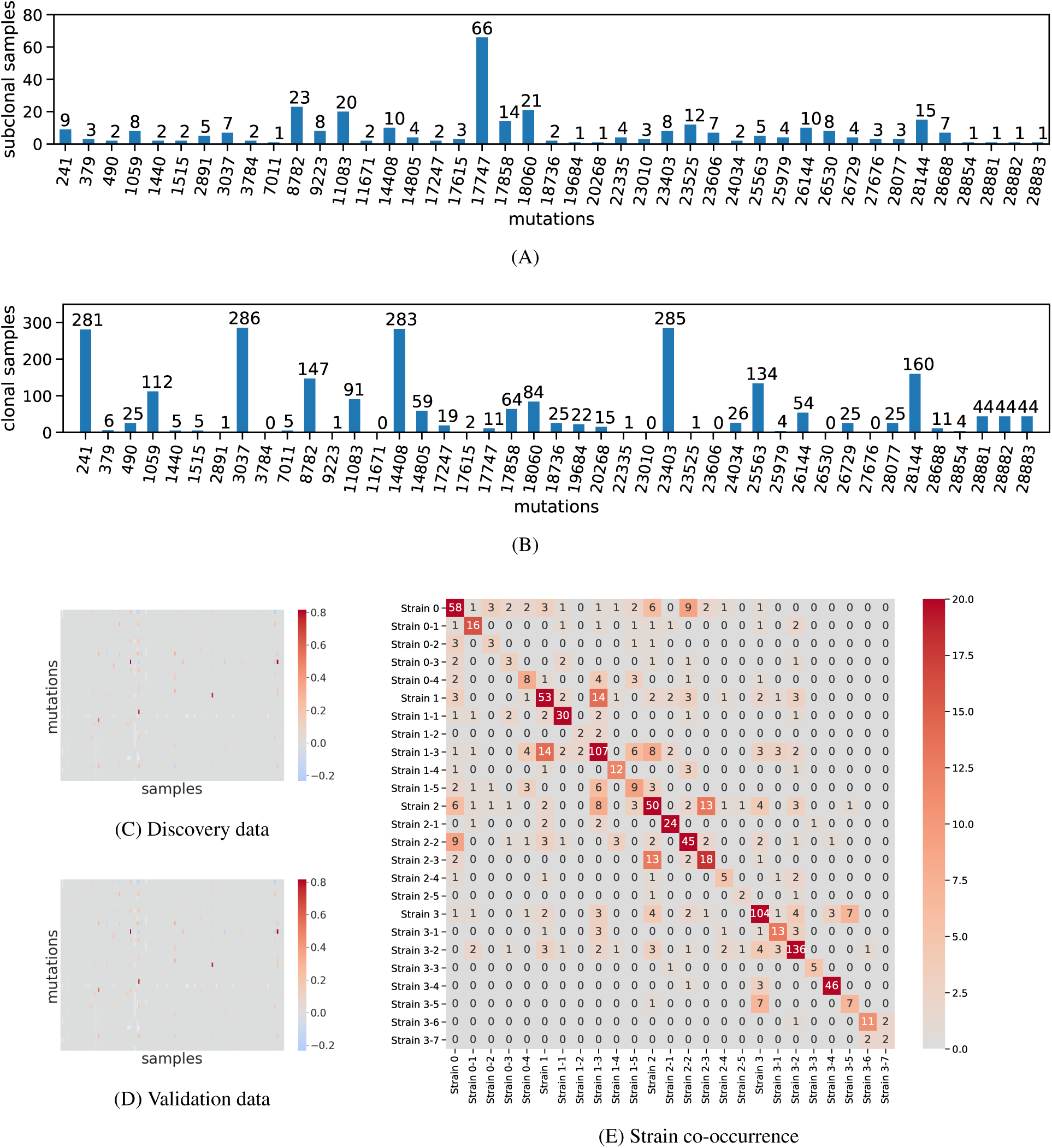
Presence of Shared Mutations and Strains across Sequence Read Archive (SRA) Samples. For each of the 43 mutations in our inferred solution on the discovery dataset, we show (A) the number of samples in which that mutation is subclonal and (B) the number of samples in which that mutation is clonal. (C) shows the reconstruction error (*F* −*BU*) heatmap for the discovery dataset and (D) the reconstruction error heatmap for the validation dataset. (E) A heatmap showing the co-occurrence of the strains in the 621 SRA samples based on the exposure inference solution. Related to Figure 4.

**Table S2:** Identified Mutations and their Phylogenetic Clades for the Tree shown in Figure S2. For each mutation, we list its gene name, protein name, amino acid change (AA), structure availability, and clade membership. The “Struct. avail.” column indicates the sources of the protein structures we collected: proteins that have solved structures in PDB are labeled with “solved” and the PDB IDs are provided; others proteins are labeled with “homology” if high-confidence homology models can be built; predicted structures are collected from folding algorithms for the remaining proteins, which are labeled with “folding” (STAR Methods); “-” means the mutation is either in a noncoding region or a synonymous/nonsense mutation, or there is no available structure that models the mutated position. Related to Table 1.

| SNV | Gene | Protein | AA | Struct. avail. | Clade 0 | Clade 1 | Clade 2 | Clade 3 |
| --- | --- | --- | --- | --- | --- | --- | --- | --- |
| C241T | noncoding | - | - | - | - | - | - | ✓ |
| C379A | ORF1a | nsp1 | V38V | - | - | - | - | ✓ |
| T490A | ORF1a | nsp1 | D75E | homology | - | ✓ | - | - |
| C1059T | ORF1a | nsp2 | T85I | homology | - | - | - | ✓ |
| G1440A | ORF1a | nsp2 | G212D | homology | ✓ | - | - | - |
| A1515G | ORF1a | nsp2 | H237R | homology | - | - | ✓ | - |
| G2891A | ORF1a | PL-PRO | A58T | homology | ✓ | - | - | - |
| C3037T | ORF1a | PL-PRO | F106F | - | - | - | - | ✓ |
| C3784T | ORF1a | PL-PRO | V355V | - | - | - | - | ✓ |
| C7011T | ORF1a | PL-PRO | A1431V | folding | ✓ | - | - | - |
| C8782T | ORF1a | nsp4 | S76S | - | - | ✓ | - | - |
| C9223T | ORF1a | nsp4 | H223H | - | - | - | ✓ | - |
| G11083T | ORF1a | nsp6 | L37F | folding | - | - | ✓ | - |
| C11671T | ORF1a | nsp6 | R233R | - | - | - | ✓ | - |
| C14408T | ORF1b | nsp12 | P323L | solved (PDB:7BV1) | - | - | - | ✓ |
| C14805T | ORF1b | nsp12 | Y455Y | - | - | ✓ | ✓ | - |
| T17247C | ORF1b | nsp13 | R337R | - | - | - | ✓ | - |
| A17615G | ORF1b | nsp13 | K460R | homology | - | - | - | ✓ |
| C17747T | ORF1b | nsp13 | P504L | homology | - | ✓ | - | - |
| A17858G | ORF1b | nsp13 | Y541C | homology | - | ✓ | - | - |
| C18060T | ORF1b | nsp14 | L7L | - | - | ✓ | - | - |
| T18736C | ORF1b | nsp14 | F233L | homology | - | ✓ | - | - |
| G19684T | ORF1b | nsp15 | V22L | solved (PDB:6VWW) | - | ✓ | - | - |
| A20268G | ORF1b | nsp15 | L216L | - | - | - | - | ✓ |
| G22335T | S | S | W258L | folding | - | - | ✓ | - |
| T23010C | S | S | V483A | folding | - | ✓ | - | - |
| A23403G | S | S | D614G | solved (PDB:6VXX) | - | - | - | ✓ |
| C23525T | S | S | H655Y | solved (PDB:6VXX) | - | ✓ | - | - |
| C23606T | S | S | R682W | - | ✓ | - | - | - |
| C24034T | S | S | N824N | - | - | ✓ | - | - |
| G25563T | ORF3a | ORF3a | Q57H | folding | - | - | - | ✓ |
| G25979T | ORF3a | ORF3a | G196V | folding | - | ✓ | - | - |
| G26144T | ORF3a | ORF3a | G251V | folding | - | - | ✓ | - |
| A26530G | M | M | D3G | folding | - | - | - | ✓ |
| T26729C | M | M | A69A | - | - | ✓ | - | - |
| G27676T | ORF7a | ORF7a | E95* | - | ✓ | - | - | - |
| G28077C | ORF8 | ORF8 | V62L | folding | - | ✓ | - | - |
| T28144C | ORF8 | ORF8 | L84S | folding | - | ✓ | - | - |
| T28688C | N | N | L139L | - | - | - | ✓ | - |
| C28854T | N | N | S194L | folding | - | - | - | ✓ |
| G28881A | N | N | R203K | folding | - | - | - | ✓ |
| G28882A | N | N | R203R | - | - | - | - | ✓ |
| G28883C | N | N | G204R | folding | - | - | - | ✓ |

